# Designing efficient randstrobes for sequence similarity analyses

**DOI:** 10.1101/2023.10.11.561924

**Authors:** Moein Karami, Aryan Soltani Mohammadi, Marcel Martin, Barış Ekim, Wei Shen, Lidong Guo, Mengyang Xu, Giulio Ermanno Pibiri, Rob Patro, Kristoffer Sahlin

## Abstract

Substrings of length *k*, commonly referred to as *k*-mers, play a vital role in sequence analysis, reducing the search space by providing anchors between queries and references. However, *k*-mers are limited to exact matches between sequences. This has led to alternative constructs, such as spaced *k*-mers, that can match across substitutions. We recently introduced a class of new constructs, *strobemers*, that can match across substitutions and smaller insertions and deletions. *Randstrobes*, the most sensitive strobemer proposed in [18], has been incorporated into several bioinformatics applications such as read classification, short read mapping, and read overlap detection. Randstrobes are constructed by linking together *k*-mers in a pseudo-random fashion and depend on a hash function, a *link function*, and a comparator for their construction. Recently, we showed that the more random this linking appears (measured in entropy), the more efficient the seeds for sequence similarity analysis. The level of pseudo-randomness will depend on the hashing, linking, and comparison operators. However, no study has investigated the efficacy of the underlying operators to produce randstrobes.

In this study, we propose several new construction methods. One of our proposed methods is based on a Binary Search Tree (BST), which lowers the time complexity and practical runtime to other methods for some parametrizations. To our knowledge, we are also the first to describe and study the types of biases that occur during construction. We designed three metrics to measure the bias. Using these new evaluation metrics, we uncovered biases and limitations in previous methods and showed that our proposed methods have favorable speed and sampling uniformity to previously proposed methods. Lastly, guided by our results, we change the seed construction in strobealign, a short-read mapper, and find that the results change substantially. Also, we suggest combining the two versions to improve accuracy for the shortest reads in our evaluated datasets. Our evaluation highlights sampling biases that can occur and provides guidance on which operators to use when implementing randstrobes.

## 1 Introduction

In sequence analyses, *k*-mers play an important role in various algorithms and approaches. For example, *k*-mers can be used as *seeds* for sequence similarity search, where a seed shared between two sequences acts as an *anchor* in order to identify similar regions between, e.g., DNA, RNA, or protein sequences. When used as seeds, *k*-mers enable rapid identification of shared regions and are used in a large number of short and long-read mapping algorithms [4,21], and other approaches for querying large sequence datasets [13].

Both a feature and a limitation with using *k*-mers as seeds is that sequences must be identical for the seed to match. In biological data, it is common that mutations in DNA occur in the form of substituted, deleted, and inserted nucleotides. In addition, common DNA and RNA sequencing techniques are noisy and introduce additional altering of the nucleic acids. In order to provide anchors also in regions with high divergence, seeds are allowed to *anchor* over mutations. *k*-mer alternatives have therefore been explored extensively in the literature, such as spaced *k*-mers [11]. See [21] for an overview of several other seeding constructs used in read mapping.

### 1.1 Strobemers

Recently, we introduced a new class of seed constructs, called *strobemers* [18]. Strobemers allow a pair of seeds to match across substitutions, insertions and deletions. Strobemers expand on the ideas of neighboring minimizer pairs [5,22] and *k*-min-mers [7] that link together neighboring minimizers [17] into a seed. At a high level, strobemers generalize this linking by considering several downstream *k*-mers within a window as potential candidates to link. Three different methods to link the *k*-mers (minstrobes, randstrobes, and hybridstrobes) were described in, [18] where the most effective seed was randstrobes. While there are applications that use other strobemer types [8], randstrobes have been most frequently used, e.g., for short-read mapping [20], transcriptomic long-read normalization [15], and read classification [23] in bioinformatic applications. Our recent proof-of-concept study also shows that randstrobes can provide accurate sequence similarity ranking through estimating the Jaccard distance [12].

In [12], we also found that the *sensitivity* of strobemers, measured as producing at least one seed match in a mutated region of fixed length, is strongly correlated with the *pseudo-randomness* of the seed construct (measured through entropy), where higher entropy yields higher sensitivity. In [12], we also introduced new strobemer variations, further improving sequence matching performance. Despite the introduction of these new variations, randstrobes remain the simplest and most used construct. Since randstrobes are now employed in multiple applications and the possibility of future applications exists, it is important to study how to construct them best.

Constructing randstrobes consists of converting strings to integers through a hash function and selecting candidate *k*-mers to link through a link function and a comparator operator (detailed definitions in Sec. 2, Methods). Randstrobes are pseudo-random seeds, meaning the linking pattern is fixed but appears random. Biases in these functions and operators through correlation will result in biases in linking the *k*-mers, making the seed less efficient for, e.g., sequence matching. Fig. 1 shows some of the sampling biases we observed in this study with different methods. So far, no evaluation has been performed of the underlying operators to produce randstrobes.

**Fig. 1:**
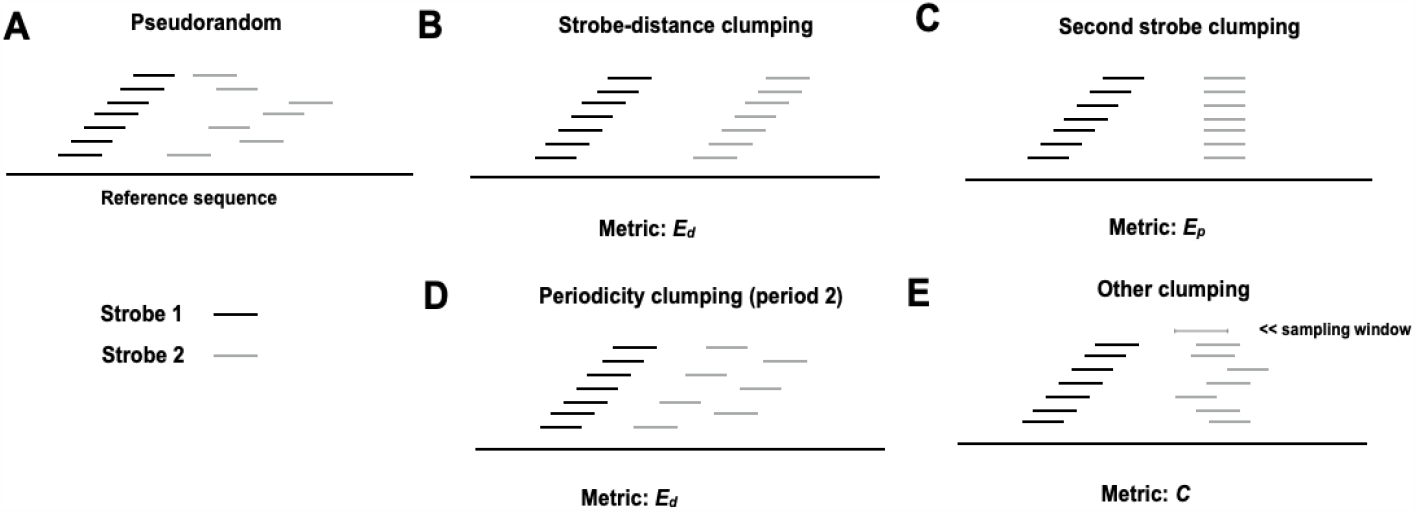
Illustration of desired random sampling of the second strobe for strobemers consisting of two strobes (case A). Whenever a pseudo-random method is used to select the downstream strobe based on the first strobe, it generates some sampling bias. Case B to E show different biases we observed in the sampling. The metrics we propose to measure the bias is displayed under each of the illustrations of cases B to E.

### 1.2 Our contribution

As randomness is important for sensitivity [12], we propose several new methods to perform the core operations in randstrobes (hashing, linking, and comparison) beyond previously published methods [18,20,23]. We also observe several types of bias (Fig. 1) with previously proposed methods and design metrics that are suitable to detect and measure them. Using new evaluation metrics, we uncovered biases and limitations in previous techniques and improved existing method to be faster and better randomly distributed. Our benchmarking of our proposed methods show that some combinations of hash functions, link- and comparison operators produce strictly favorable results to previously proposed methods, improving e.g., seed uniqueness, sampling uniformity, and construction runtime.

In addition, one of the randstrobe construction methods we present is based on a Binary Search Tree (BST) that lowers the time complexity of constructing randstrobes from what was previously reported in [18]. The method is much faster than other methods when the strobe selection window is large and achieve comparable randomness to the best performing method for large windows. As runtime is important in several time consuming bioinformatics applications, such a BST implementation may be useful for applications where a large sampling window is desired.

Finally, we find that the combination of link function and comparator used in the short-read mapper strobealign [20] perform strictly worse regarding seed uniqueness than other methods. Guided by this observation, we changed the seed construction strobealign. While our new implementation does not increase *strobealign’s* accuracy on our benchmarked datasets, we observe that the accuracy improves substantially for an approach that selects the best alignment score per read from our modified version and the default version of strobealign. This finding can be used to increase strobealign’s accuracy further. In summary, our evaluation highlights linking biases that can occur and provide guidance for which operators to use when implementing randstrobes.

## 2 Methods

### 2.1 Definitions

We use 0-indexed notation. We typically use *S* and *T* to denote strings, and we use the notation *S*[*i* : *j*], *i < j* to refer to a substring starting at position *i* and ending and including the character at position *j* in *S*. We let the | · | operator denote the length of strings. Here, our alphabet consists of the letters (or *nucleotides*) (Σ = *A, G, C, T*). We use *h*(*x*) → *z*, where *x* and *z* are integers to denote a hash function without specifying the underlying function. As for representation in memory, DNA strings shorter or equal to 32 nucleotides can be stored with 64-bit integers by encoding A, C, G, and T as 00, 01, 10, and 11, respectively. Other letters, such as N for unknown nucleotide, will be ignored.

In this study, we will consider strings shorter than 32 nucleotides and use variable *x* to represent the 64-bit integer values encoding either the 2-bit representation of the string or the hash value resulting from hashing the string. While the constraint on 32 nucleotides is often seen as a limitation for *k*-mer-based methods in bioinformatics, it does not represent a limitation to the same extent for strobemers as they consist of several combined *k*-mers, which we discuss in Section 2.2. Finally, we use & for bitwise AND, ⊕ for bitwise XOR, ‖ for concatenation (e.g., concatenating two 64-bit integers into a 128-bit representation), and % for the modulo operator. We also use *B*(*x*) to represent the function that returns the number of set bits in *x*.

### 2.2 An overview of constructing strobemers

A *k*-mer is a substring of *k* nucleotides in a biological sequence *S*. Consequently, a *k*-mer only needs the length of the substring, *k*, as a parameter to be specified. A strobemer is a set of linked *k*-mers. Specifically, a strobemer consist of *n l*-mers *l*_0_, …, *l*_*n*−1_, denoted *strobes*, where the first strobe *l*_0_ has a determined position *i* in *S*. Downstream strobe *l*_*m*_, *m* ∈ [1, *n* − 1] is selected in an interval *S*[*i* + *w*_*min*_ +(*m* − 1)*w*_*max*_ : *i* + *mw*_*max*_] in *S*, and *linked* (appending the strobe to previous strobes) to the *m* previous strobes. Here, *w*_*min*_ and *w*_*max*_specify the range of the sampling window. For example, strobe *l*_1_ is sampled in *S*[*i* + *w*_*min*_ : *i* + *w*_*max*_] and linked to *l*_0_.

Since we consider 64-bit integer representations of the strobes in this study, we will from now on refer to the strobes as *x*_0_, *x*_1_, … *x*_*n*−1_ and, when clear from context, we alternate *x* to mean either the strobe itself or its integer representation. This is also the reason we use the more general term *linking* instead of *appending* (strobes to the seed), as the linking method will vary with the strobe representation, as we discuss in detail in the next section.

The methods to select strobes differ [18], and using alternating strobe lengths has also been explored [12]. However, randstrobes were shown to be more sensitive for sequence matching than other methods using fixed strobe lengths (minstrobes and hybridstrobes) [18], and simpler to construct than alternating strobe lengths (altstrobes and multistrobes) [12], and is so far most commonly implemented in practice, e.g., [20,15,23]. Therefore, we will consider only the randstrobes method in this study. Randstrobes are parameterized by (*n, l, w*_*min*_, *w*_*max*_). The novelty compared to, e.g., *k*-mers and spaced *k*-mers is that strobemers allow flexibility in the strobes’ spacing and can produce matches between two sequences in a region with insertions or deletions.

### 2.3 Strobemer construction: constraints and objectives

Let 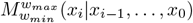, or simply *M* when context is clear, be a method to sample a strobe *x*_*i*_ in a window given by its parametrization (*n, l, w*_*min*_, *w*_*max*_). We put the following constraints on *M*.

**Constraint 1:** *M* selects *x*_*i*_ based on, and only on, the sequence information of *x*_*i*−1_, …, *x*_0_.

**Constraint 2:** *M* is deterministic. That is, for two identical strings *S* and *T*, the same strobes are produced.

We want to find a method *M* such that

**Objective 1:** Maximize *H*(*M* (*x*_*i*_|*x*_*i*−1_, …, *x*_0_)), where *H* denotes the entropy. Intuitively, *M* should sample *x*_*i*_ as uniform within the window as possible, regardless of previous strobes and the sequence in the window.

**Objective 2:** *M* constructs randstrobes as fast as possible.

Constraint 1 is necessary to rule out degenerate but high entropy solutions for sequence matching. An example of a degenerate approach is using a (pseudo) random number generator (RNG) such as `rand()` in C++. Such a method has good entropy. However, assume two similar strings, *S* and *T*, where one string contains a deletion. The RNG will (in all likelihood) produce different numbers as soon as the deletion is encountered and will continue to produce different numbers throughout the remaining part of the string. Such an approach cannot be used for string-matching applications. Therefore, the method’s decision must be based on, and only on, the underlying sequence. As for the objectives, Objective 1 describes a conditional entropy (conditioned on the first strobe and the window), which is challenging to measure. We cannot only evaluate entropy by measuring the uniformity of sampling sites within a window across a sequence. For example, assume a method that selects a strobe if it is identical to the previous strobe (otherwise, it uses some other decision). If the distance between two consecutive identical strobes is uniformly distributed, the method will appear to have a perfect entropy while it, in fact, has low entropy for other input. Related, it is easy to display high entropy for randomly generated sequences. However, we are primarily interested in what happens in repetitive regions, common in biological sequences, and more challenging to produce sampling uniformity over. Objective 2 is straightforward.

### 2.4 Constructing randstrobes

The process of creating randstrobes can be separated into four modular components: 1) Hashing the strobes, 2) linking the strobes, 3) the use of sampling comparator when linking, and 4) the construction of the final seed hash value. We discuss each of the components below and suggest different functions to perform them.

#### Hashing strobes

Since each strobe is represented as a 64-bit integer using the binary encoding, the integers can further be hashed. The reason for hashing a strobe *x* as *z* = *h*(*x*) is that it can improve the pseudo-randomness. We evaluate the following hash functions for the strobes.

1. *h*_NO_(*x*): The original 2-bit encoding of nucleotides is used without applying a hash function.
2. *h*_TW_(*x*): *Thomas Wang hash* [1], an invertible hash function used, e.g., in the popular aligner minimap2 [10].
3. *h*_XX_(*x*): *XXHash* [3].
4. *h*_WY_(*x*): *WYHash* [2].

Previously, *h*_NO_(*x*) was used in [18] and *h*_TW_(*x*) was used [20]. This is the first study using *h*_XX_(*x*) and *h*_WY_(*x*) as hash functions to construct randstrobes.

**Linking strobes** The second strobe *x*_1_ is *linked* to the first strobe *x*_0_ by selecting the candidate strobe 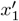 in the window that minimizes or maximizes the link function ℓ. For example, in the first strobemers study [18], two link functions were used. The first was 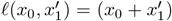 mod *p, p* ∈ *Z* (originally proposed in the preprint [19]). The second one was 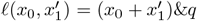, where *q* is a bitmask of 16 ones’ on the lowest significant bits and remaining 0’s (proposed as faster alternative in the final publication [18]). We call these function ℓ_MOD_ and ℓ_AND_, respectively. Furthermore, two additional link functions were described in [20,23] that we denote ℓ_BC_ and ℓ_XOR_. Here we propose three more alternatives: ℓ_XV_, ℓ_CC_, and ℓ_MAMD_. We provide formal definitions of all the link functions below.

– ℓ_MOD_(*x*_0_, *x*_1_) = (*h*(*x*_0_) + *h*(*x*_1_)) mod *p, p* ∈ *N*. (See [18])
– ℓ_AND_(*x*_0_, *x*_1_) = (*h*(*x*_0_) + *h*(*x*_1_))*&q, q* ∈ *N*. (See [18])
– ℓ_BC_(*x*_0_, *x*_1_) = *B*(*h*(*x*_0_) ⊕ *h*(*x*_1_)). (See [20])
– ℓ_XOR_(*x*_0_, *x*_1_) = *h*(*x*_0_) ⊕ *h*(*x*_1_). (See [23])
– ℓ_XV_(*x*_0_, *x*_1_) = *h*(*x*_0_ ⊕ *x*_1_). (Proposed in this study)
– ℓ_CC_(*x*_0_, *x*_1_) = *h*(*x*_0_ ‖ *x*_1_). (Described in the pseudo code in [18] but never studied)
– ℓ_MAMD_(*x*_0_, *x*_1_) = (*h*(*x*_0_) mod *p*) + (*h*(*x*_1_) mod *p*) mod *p, p* ∈ *N*. Similar to ℓ_MOD_ but uses a BST. (Proposed in this study)

The ℓ_MAMD_ and ℓ_MOD_ are theoretically identical. However, in practice the ℓ_MOD_ can overflow the integer limit (2^64^ − 1), while ℓ_MAMD_ can not if *p <* 2^63^ − 1. Furthermore, ℓ_MAMD_ uses a BST (ℓ_MAMD_ is described in detail in Supplementary Section A1) with a different computational complexity. We will discuss the computational complexity of all methods at the end of Section 2.3. In this section, we only discussed linking the first two strobes. Linking additional strobes can be done recursively by applying the same link function between the previous resulting randstrobe hash value *b* with the next candidate downstream strobes *x*_*m*_, *m >* 2 as ℓ(*b, x*_*m*_).

#### Sampling comparator

The sampling comparator is closely tied to the link-function. Specifically, the comparator function, here denoted *c*(·), specifies the criteria for which we select strobe *x*_1_ among candidates 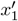. To our knowledge, the only sampling comparator that has been proposed is 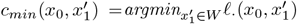 [18,20,23], where *W* is the collection of strobes in the window defined by *w*_*min*_ and *w*_*max*_. In this study we propose 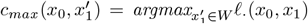. Another example of a comparator operator could be, e.g., selecting the median value.

The comparator can influence the result for some hash and link constructions. A concrete example is constructing randstrobes by using 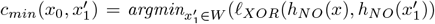. Consider a repetitive region consisting of the same short repeat with occasional variations (e.g., a telomeric region). Our method will be biased towards selecting identical strobes as identical values in *x* and *x*_1_ will have an XOR value of 0. Here, a *max* comparator will likely sample strobes with variants in such region, thus improving the uniqueness of seeds.

#### The final seed hash value

We have so far discussed only how to select strobes. However, once the strobes have been decided, we need to represent the randstrobe with a *final hash value*. The final hash value is what should be indexed and queried, for e.g., a seed-and-extend mapping framework. We denote the function to produce the final seed hash value as *f* (*x*_0_, …, *x*_*n*_). We need the function *f* to be as uncorrelated with the link-function as possible. If we would use the hash value that comes out of ℓ(*x*_0_, *x*_1_), with, e.g., *c*_*min*_, we are projecting hash values to the minimum value in each window. This leads to unnecessary hash collisions compared to a uniform hash function. Furthermore, as mentioned in [18], it is important to avoid symmetric functions *f* (*x*_0_, *x*_1_) = *f* (*x*_1_, *x*_0_) (e.g, *f* (*x*_0_, *x*_1_) = *x*_0_ + *x*_1_) if distinguishing direction from, e.g., inversions is important (although a symmetric function is used to forward and reverse complements seeds in, e.g., read mapping [20]. Taking into consideration the above we use

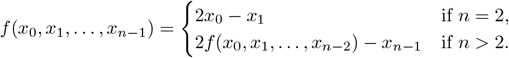

This formulation allows *f* not to have any apparent correlation with any of the benchmarked link-functions, as we will see in the results.

#### Linking more than two strobes

Generally, to link *x*_*m*_, to *x*_0_, … *x*_*m*−1_, *m* ∈ [2, *n* − 1], we use 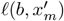, where 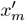 are the candidate strobes in the window, and *b* denotes a *base value* calculated from the previous *m* strobes. We set the *b* equal to the previous strobes’ final hash value, e.g., *b* = *f* (*x*_0_, *x*_1_) and 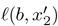 in the case of three strobes. This method can be applied recursively.

#### Time complexity of randstrobe construction

Before discussing computational complexity we make the following classifications of our link functions:

1. **Cheap computation**: This group includes ℓ_MOD_, ℓ_AND_, ℓ_BC_, ℓ_XOR_ and ℓ_MAMD_. We denote them as computationally cheap because the hashing and linking can be separated. That is, we only need to calculate hash values once for each strobe, and the link function can be applied after.
2. **Expensive computation**: This group includes ℓ_CC_, and ℓ_XV_. For these methods we need to evaluate the hash value for the combination of *x*_0_ and all its candidate downstream strobes, for each new *x*_0_.

The time complexity of constructing randstrobes from a string of length |*S*| varies with the link-function class. Let *t*_*h*_ be the time complexity for the hash function, *n* the number of strobes, and *W* = *w*_*max*_ −*w*_*min*_ +1 be the window size. Then, *S* − *nw*_*max*_ *l* + 1 the number of randstrobes constructed from *S*. We assume that the linking operators (i.e., +, &, ⊕, mod, ‖) can be performed in constant (*O*(1)) time, although the runtimes vary among the operators with ⊕; being cheaper to perform while ‖ being relatively expensive.

Expensive computation methods perform (1 + *nW*) hash calculations, and *nW* other operations (such as +, &, ⊕, mod, ‖), per randstrobe. So the total complexity is *O*((|*S*| − *nw*_*max*_ − *l* + 1)((1 + *nW*) *t*_*h*_ + *nW*)). Cheap computation methods spend at most (|*S*| − *l* + 1) hash calculations and (|*S*| − *nw*_*max*_ − *l* + 1)(*nW*) on other operations, in total. So the total complexity is *O*((|*S*| − *l* + 1)*t*_*h*_ + (|*S*| − *nw*_*max*_ − *l* + 1)(*nW*)). If we assume that *S >> nw*_*max*_ *l* + 1 and *t*_*h*_ = *Ω*(1) (i.e., the complexity of *t*_*h*_ is at least a constant), we can simplify the expression of the time complexity of expensive computation methods and cheap computation methods to *O*(|*S*|*nWt*_*h*_), and *O*(|*S*|*t*_*h*_ + |*S*|*nW*), respectively.

Lastly, the ℓ_MAMD_ link function is part of the cheap computation category. However, the time complexity is further reduced to *O* (|*S*|*t*_*h*_ + |*S*| *n* log *W*) through the logarithmic time complexity of searching for elements (see Supplementary Section A1 for details). While the BST implementation increases the constant coefficient through the BST overhead, we will see that the speed-up is substantial for large windows. We have abstracted over the exact time complexity of the hash functions. The cheapest computation is *h*_*NO*_ which only streams over the sequence without performing hashing. Some hash functions also support streaming [14] and can lower *t*_*h*_.

### 2.5 Evaluation Metrics

There are different sampling biases that can arise as illustrated in Fig 1. We were not able to find a singular metric that captured all of these biases, instead we propose four suitable metrics that would capture cases B-E in Fig 1. A desirable result is that the selection of the second (or any downstream) strobe is performed as uniformly in the window and as independently of previous seed as possible. Several seed-based applications also requires fast construction; therfore, we also benchmark construction runtime.

#### Notation for evaluation metrics

Let *N* be the total number of seeds constructed from a string *S*, and *M* the number of seeds with distinct final seed hash value in *S*. Recall that (*n, l, w*_*min*_, *w*_*max*_) parameterize the number of strobes, their length, and the minimum and maximum window offset for the sampling window of the randstrobes. We let *i* and *j* be index variables over the set of randstrobes seeds sorted by their first strobe position. Since we here sample one randstrobe per position in *S*, the index variables are equivalent to the start position of the randstrobe seed on *S*, and the *N* seeds can be ordered with respect tp the start position on *S*. We let *s*_*ik*_ refer to the *k*th strobe in seed *i* and *p*_*ik*_ to its position in *S*.

#### E-hits

The E-hits metric was introduced in [20]. It provides a number between 1 and |*S*|, which is the expected number of times a seed occurs in the reference. The E-hits metric was used as a measure for expected seed repetitiveness in *S* when sampling reads uniformly at random from a reference string *S*, assuming *S* is much larger than the span of the seed [20]. We restate the E-hits metric here for self-containment. Let *i* ∈ [1, *M*] be an index variable over the set of distinct seeds in *S* and *N > M* be the total number of seeds in *S* (multiset). Let *x*_*i*_ denote the number of times seed *i* occurs in *S*. Let *q*_*i*_ be the probability of producing seed *i* when selecting a seed randomly from the *N* seeds. The E-hits metric is then the expected value over seed hits *E*[*X*] computed as

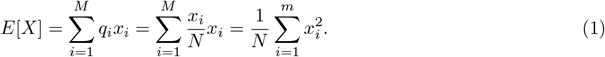

In this study, seeds are represented as hash values. The above formula is equivalent if we replace the notion of a seed with the hash value representation of a seed. In this case, E-hits measures the expected number of identical hash values, which includes both repetitive seeds and non-desired hash collisions. We will measure the E-hits for the final seed hash values produced with *f*, and denote this quantity *E*_*f*_. This is the same use of E-hits as in [20].

#### E-hits of inter-strobe distance and strobe position

The idea and formulation of E-hits can be used to measure the repetitiveness of other quantities. To measure biases B and D in Fig 1, we look at the distribution of inter-strobe distances within a randstrobe. Let *d*_*jk*_ be the distance between the first strobe and the *k*th strobe in seed *j* (in this study, we only consider *k* = 2 or *k* = 3). We can then let *x*_*i*_ in Eq. 1 instead denote the number of times we observe distance *d*_*jk*_. The E-hits formula then measures the expected number of times we observe the distance *d*_*jk*_ when randomly drawing a seed from *S*. We denote this quantity as *E*_*d*_, we omit index variable *k* when it is clear from the context.

We measure bias C by computing the repetitiveness of the position of *k*th strobes in *S*. Along the same vein as *E*_*d*_, we let *x*_*i*_ in Eq. 1 instead represent the number of times we observe the *k*th strobe in any randstrobe being at position *p* in *S*. For this quantity, the E-hits formula then measures the expected number of times position *p* was sampled as the *k*th strobe when drawing a seed uniformly at random from *S*. Similarly to *E*_*d*_, we denote this quantity as *E*_*p*_ and omit index variable *k* when it is clear from the context. We can compare *E*_*d*_ and *E*_*p*_ against a reference method that generates strobe positions uniformly at random within the sampling windows. This comparison allows us to study the relative magnitudes of the bias between the methods.

#### The conflict metric

To study which complex dependencies as depicted in Case E in Fig 1, we introduce the *conflict metric*, which aims to measure the size of the overlaps of strobes from a set of neighbouring randstrobes with start positions in [*i, j*], *i < j*. An overlap higher than what is expected under random sampling indicates selection bias. Let *o*(*i, j, k*) = *max*(0, *l* − |*p*_*jk*_ − *p*_*ik*_|) measuring the number of overlapping positions of the *k*th strobe between two randstrobes *i* and *j*. Then 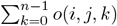 is the total number of overlapping positions between two randstrobes. The conflict metric for randstrobe *i* is then defined as

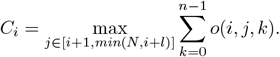

In other words, *C*_*i*_ is the largest observed overlap with any of the *l* consecutive downstream randstrobe seeds. We let the conflict metric (*C*) be the value of *C*_*i*_ averaged over all seeds in *S*. As with the other metrics, we can compare *C* against a reference method generating strobe positions uniformly at random within the sampling windows. In addition to measuring bias E, this measure also captures biases B, C, and D (Fig 1).

The above formula does not take into account that strobes of different orders (*k*) between neighboring randstrobes might overlap. However, even if this is possible for some values of *w*_*min*_, it does not originate from the bias that we want to measure, and can therefore be omitted.

## 3 Results

We evaluated all compatible combinations of ℓ, *c* and *h*. Some hash functions and link-functions that are incompatible such as using *h*_*TW*_ and ℓ_*CC*_ with strobes larger than 16nt (32 bits) because *h*_*TW*_ is designed for 64-bit integers. We used *p* = 100, 001 for ℓ_*MAMD*_ and ℓ_*MOD*_ in our experiments. The evaluation is available at https://github.com/Moein-Karami/RandStrobes/.

### 3.1 Experiment setup

As discussed in section 2.3, it is easy to produce randstrobes with high entropy if the underlying sequence in [*w*_*min*_, *w*_*max*_] has high entropy (e.g., randomly generating letters in {A,C,G,T}). Therefore, we are interested in evaluating pseudo randomness in repetitive regions, common in biological sequences. We use both a simulated highly repetitive sequence, as well as chromosome Y from the CHM13 human assembly [16], including telomere regions. We simulated a repetitive sequence *S* as follows. A sequence *T* consisting of 25 nucleotides A, G, C, and T was randomly generated and appended to *S*. We then simulated a new copy of *T′* from *T* by mutating each position in *T* with a probability of *p* = 0.02, where the mutation could either be a substitution, insertion, or deletion with equal probabilities. *T′* was then appended to *S* and used as the new template to simulate the next copy *T″*. This recursive procedure was repeated 40,000 times. If the length of any template copy decreased to below 15 nucleotides, we only considered substitutions and insertions for those templates. This process resulted in a string of approximately 1,5 million nucleotides.

We used randstrobe parameter value of *n* = 2, *l* = 20, *w*_*min*_ = 21, *w*_*max*_ = 100. In addition, we also report the results for three strobes (*n* = 3) in Supplementary Section A3. Finally, since the metrics we use could be difficult to interpret in a vacuum, we have, when applicable, also included suitable reference values. These reference values could either be *k*-mers with size *k* = *nl*, or a fully random method, denoted *uniform*, that produces randstrobes by uniformly at random selecting a position in the sampling window [*w*_*min*_, *w*_*max*_] (using `rand()` in C++). We remark that method produces different randstrobes from the same sequence. Thus, we cannot use uniform randstrobes for anything other than providing best-case reference values for other methods.

To evaluate runtime, we use an *E. coli* genome (strain NZ CP018237.1) of roughly 5.5 million nucleotides. We report the median runtime from 25 runs. We consider the difference between the starting and the finishing time of creating and storing randstrobes in a vector as the execution time. The experiments were run on an Linux computer, with an Intel i7-4510U CPU at 2.00GHz, compiled with gcc with flag `-O3`. For the runtime, we evaluated randstrobes parametrized as (*n* = 2, *l* = 20, *w*_*min*_ = 21, *w*_*max*_ = 100) and (*n* = 2, *l* = 20, *w*_*min*_ = 21, *w*_*max*_ = 1000) since the window size affects runtime. Strobemers with *n >* 3 show no substantial gain in the context of sequence matching at the cost of additional runtime [12](although they have been modified and used for specific scenarios [8]). Also, the relative performance can be extrapolated from the *n* = 2 and *n* = 3 cases, since the construction is recursive, therefore, we omit them in this study.

### 3.2 Pseudo-randomness

Overall, the expensive-computation linking methods ℓ_*CC*_ and ℓ_*XV*_ yield the most desirable pseudo-randomness across the three metrics *E*_*d*_, *E*_*p*2_, and *C* (Fig. 2A-C). When comparing the less computationally expensive methods, we observe that bias in only one or two of the metrics we designed, which motivates the analysis of pseudo-randomness using several metrics. The following sections will analyze the results when constructing randstrobes with two strobes. We see similar trends when constructing randstrobes with three strobes (discussed in Supplementary Section A2).

**Fig. 2:**
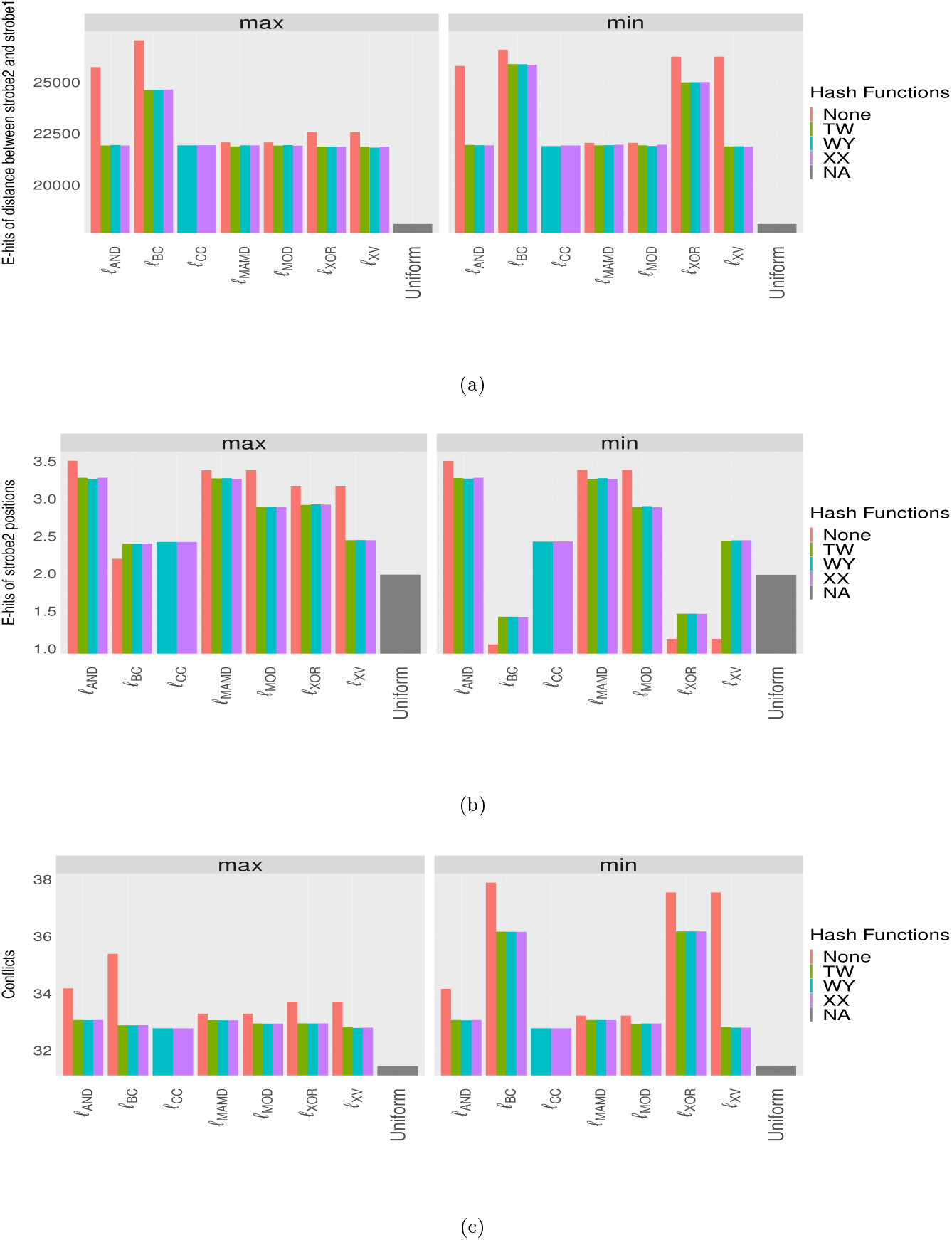
Results for metrics *E*_*d*_, *E*_*p*_, and *C* for randstrobes with parameter settings (*n* = 2, *l* = 20, *w*_*min*_ = 21, *w*_*max*_ = 100) for the repetitive sequence dataset. The *x* -axis shows the different linking methods, and the max and min comparator are shown in left and right panels, respectively. The *x* -axis also contains the reference value denoted *uniform*, indicating near perfect randomness. For metric *E*_*d*_, the lowest possible value is desirable, as the *E*_*d*_ metric reaches the minimum under a uniform distribution. For *E*_*p*_ and *C*, a value close to the uniform is desirable, which is different from the minimum, due to bias B in Fig. 1. The *y* -axis has been set to start close to the lowest observed value to better illustrate differences between the methods.

#### Bias B and D

Biases B and D (Fig. 1), characterized best with *E*_*d*_, reflect a sweeping bias in the sampling (Fig. 1). Our benchmarks are performed on a repetitive sequence, so some bias in *E*_*d*_ is expected, which is not reflected in the uniform best-case scenario. We observe that all methods have higher *E*_*d*_ than the uniform method, but ℓ_*BC*_ and ℓ_*XOR*_ have a substantially higher bias (Fig. 2a). Both of these link methods depend on the XOR operator.

#### Bias C

Bias C (Fig. 1) indicates over-sampling a given position and is best characterized with *E*_*p*_. Again, all methods deviate from the uniform (Fig. 2b). Methods such as ℓ_*AND*_, ℓ_*MAMD*_, ℓ_*MOD*_, and ℓ_*XOR*_ indicate a clear over-sampling of the same position, while ℓ_*CC*_ and ℓ_*XV*_ have values relatively near uniform. Notably ℓ_*BC*_ and ℓ_*XOR*_ with *c*_*min*_ have lower values of *E*_*p*_ than uniform. However, this is undesirable and further illustrates how these methods suffer from biases B and D, which results in lower values on *E*_*p*_ than expected. We also note a difference between ℓ_*MOD*_ and ℓ_*MAMD*_ when hashing. This difference is likely because ℓ_*MOD*_ is allowed to overflow increasing randomness, while ℓ_*MAMD*_ is not. This overflow does not happen for *h*_NO_ since the individual strobe values are always below 2^2*l*^ = 2^40^.

#### Non-trivial bias with the conflict metric

We designed the conflict metric to detect any sampling biases that are difficult to classify (Case E in Fig. 1). However, our results indicate that this metric did not pick up any notable biases that our metrics *E*_*d*_ and *E*_*p*_ did not already show. However, the conflict metric agrees with the two other metrics and and acts as a supplementary aspect to them. The min comparator performs substantially worse for ℓ_*BC*_ and ℓ_*XOR*_, as well as ℓ_*XV*_ when a hash function is not applied. elaborate on this result in the following sections.

#### Importance of using a hash function before linking

First, using a hash function before linking is performed for all linking methods except for ℓ_*CC*_, which applies the hash function after concatenation of the strobes. We will therefore exclude ℓ_*CC*_ from the discussion in this section. Our experiments show that using a hash function increases the randomness of all methods. Most notable is the difference for ℓ_*XV*_. Link functions that use the XOR operator are generally sensitive to when strobes are similar. This can be understood by considering that two identical strobes will always be projected to a value of 0. Similarly, two strobes with only a substitution difference would only have bits set where the mutation occurs unless a hash function is used. Therefore, link methods based on XOR are sensitive to the hash function used in repetitive regions. We observe that *h*_WY_, *h*_XX_, and *h*_TW_ have near identical results, except when using the ℓ_*MOD*_ link function where *h*_WY_ and *h*_XX_ are slightly preferred.

#### Max comparator is better in repetitive regions

We can see in Fig 2 that the max comparator is always either as good or better than the min comparator. The largest differences are observed for the link functions based on the XOR operator. In repetitive regions, the min comparator is likely to pick the same or similar strobes since many bits will be set to 0, while the max comparator inverts this behavior and instead is more likely to select as different strobes as possible (to increase the likelihood of significant set bits). This behavior is beneficial in repetitive regions where we benefit from more seeds with unique mutations. Our experiments indicate that the comparator affects mainly ℓ_*BC*_, ℓ_*XOR*_. However, these link functions are used in bioinformatics tools [20,23], highlighting that sampling could be improved in repetitive regions.

#### Seed repetitiveness

We have, in previous sections, investigated linking bias arising from combinations of hash, link and comparator functions. However, overall seed repetitiveness in the reference is one of the most important measures for applications such as average nucleotide identity (ANI) estimation or sequence mapping. First, we confirmed that using our final hash function *f* to represent a seed resulted in a non-visible number of hash collisions, as measured by the fraction of unique final hash values to the number of unique randstrobe seeds obtained from extracting the sequence at the sampling positions (Fig. 6). The lowest ratio of unique final hash values to actual seeds we observed was 0.9981 for ℓ_*BC*_ with *c*_*min*_. Many of the methods had no collisions (ratio equal to 1.0). *K*-mers without hashing (used as reference value) has a ratio of 0.9996. This is because we represent *k*-mers as two adjacent strobes (*x*_0_, *x*_1_) with the same final function as the randstrobes (*f* (*x*_0_, *x*_1_) = 2*x*_0_ − *x*_1_) because they do not fit into 64 bits. Regardless, this is a very small amount of collisions and should not affect the analysis.

Given that there were no significant amount of hash collisions in our seeds, we computed the E-hits of the final seed hash value (*E*_*f*_). We also included *k*-mers of length 40 in this experiment for reference. Agreeing with previous analyses, we observed that it is important to use a hash function before linking strobes and that ℓ_*BC*_ and ℓ_*XOR*_ should, if used, be combined with the max comparator (Fig. 3). Additionally, we see that randstrobes have lower *E*_*f*_ than *k*-mers for most hash and link functions, but can increase repetitiveness with some combinations if a hash function is not applied on the strobes.

**Fig. 3:**
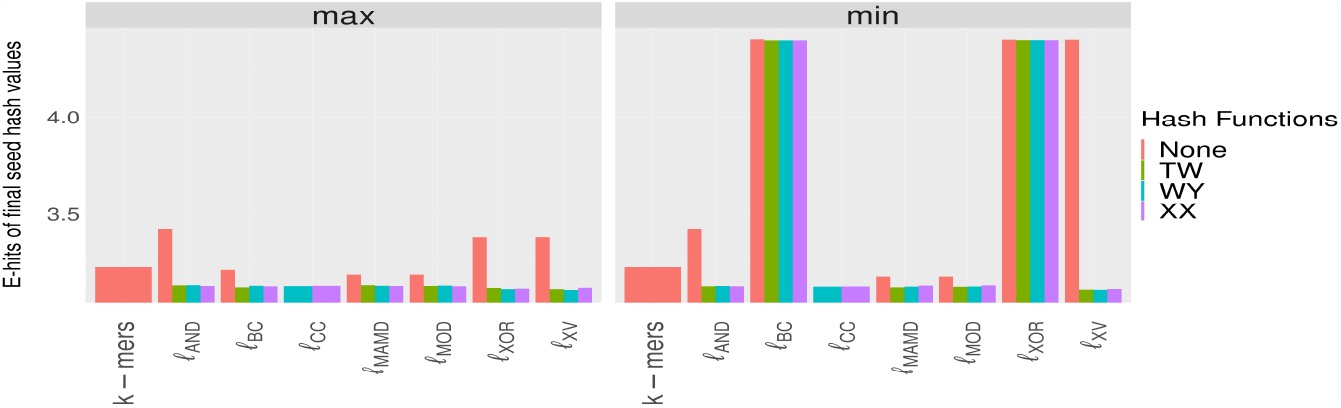
E-hits of seed hash values for various hash functions (color), link functions (x-axis) and comparators (panels) used to construct randstrobes with parameters (*n* = 2, *l* = 20, *w*_*min*_ = 21, *w*_*max*_ = 100). The x-axis also contains reference values for *k*-mers of size *k* = 40. The y-axis has been cut at 3 to better illustrate differences.

### 3.3 Runtime performance

Figure 4 shows the construction time for window sizes using *w*_*max*_ = 100 and *w*_*max*_ = 1000, respectively. Despite the expensive-computation methods (ℓ_*CC*_ and ℓ_*XV*_) are performing a factor of *nW* more hash computations, they are only about twice as slow on the smaller window (with *h*_*WY*_) to cheap computation methods (Fig. 4A) and about four times slower for large windows (Fig. 4B). One explanation could be that the window is fitting in cache, resulting in that the much higher amount of hashing calls are cheap compared to loading new parts of the array with strobes into cache. We also observe that the ℓ_*MOD*_ based on typically expensive modulo computation is substantially slower than other methods in the cheap-computation class. Finally, when constructing randstrobes with large windows, ℓ_*MAMD*_ is much faster than other methods Fig 4b. This is due to the BST implementation instead of a linear search across each window. However, due to its special updating technique utilizing arithmetic properties of the modulo operator, the method can only be used with the modulo link function. As for the hash functions, *h*_*WY*_ performs better than *h*_*XX*_ on our data for the expensive computation class, where strobes are represented by a struct of two 64-bit integer strobes. The best timing results in the expensive-computation class was ℓ_*XV*_ combined with *h*_*WY*_, albeit with a small margin to ℓ_*CC*_ combined with *h*_*WY*_.

**Fig. 4:**
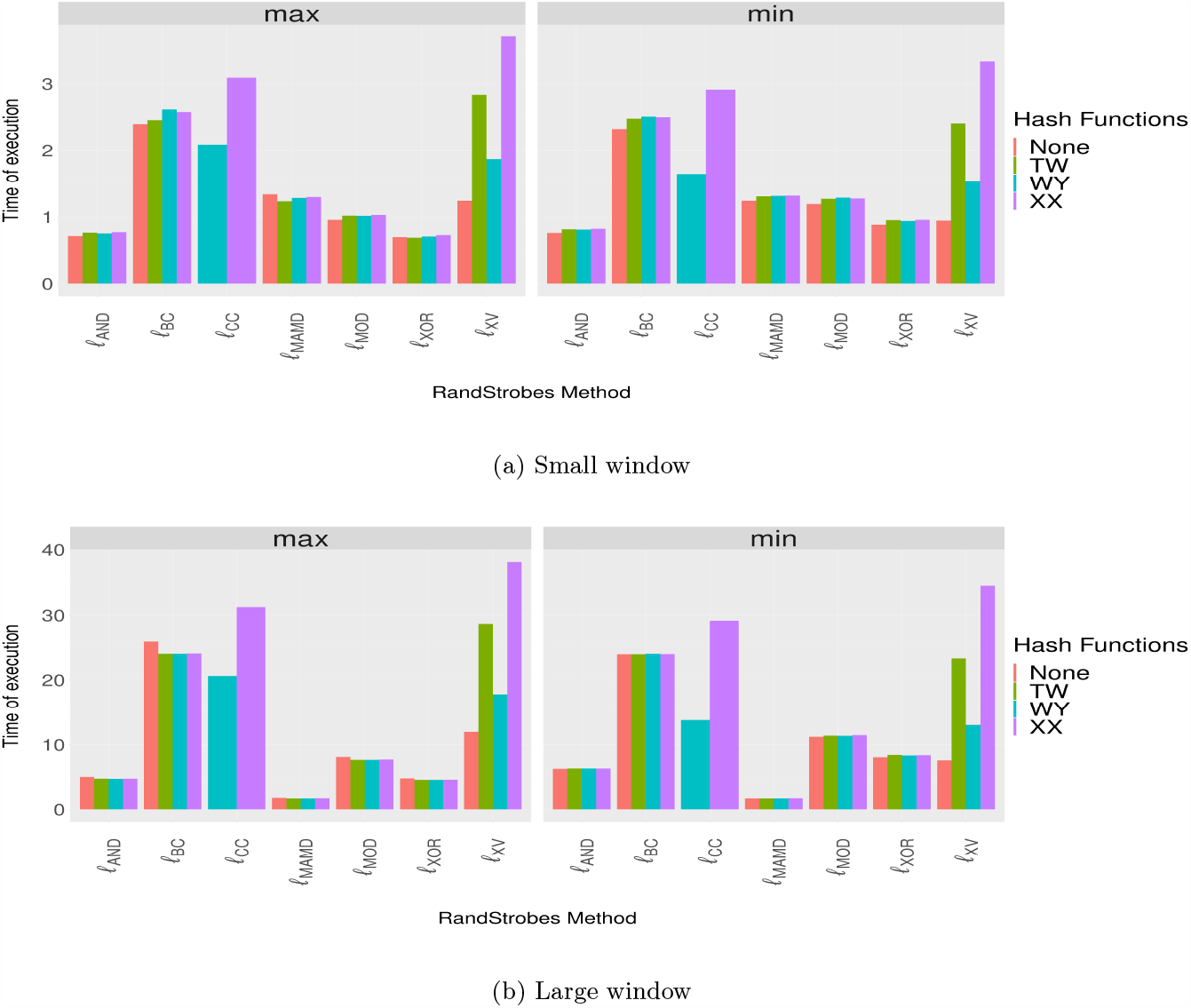
Median runtime (seconds) on 25 instances for each combination on an E. coli genome of 5.5 million nt. Each combination generated randstrobes with *n* = 2, *l* = 20, *w*_*min*_ = 21, and *w*_*max*_ = 100 (Panel A) and *w*_*max*_ = 1000 (Panel B).

### 3.4 Randstrobes in large windows

The ℓ_*MAMD*_ link function opens up the possibility to construct randstrobes in large windows. We were interested in the uniqueness of seeds that ℓ_*MAMD*_ produced compared to the best-performing method ℓ_*CC*_ (using *c*_*max*_). For this analysis we increased *p* to 19,019,684,767,739,993 as the window sizes tested were approaching previous *p* (100,001), which is not good for pseudo-randomness. We investigated the expected uniqueness (E-Hits) of the seeds computed across chromosome Y of the CHM13 assembly (Fig. 5, left panel). In the figure, a window size of 0 corresponds to *k*-mers of size 256. We make two key observations about the uniqueness of seeds. First, we note that there is no substantial difference between the two link functions on chromosome Y from the CHM13 assembly, including telomere regions and many repetitive multigene families. Second, we observe that the E-hits function is not linearly decreasing, which we initially expected. Minimum repetitiveness occurs at *w*_*max*_ = 2, 000 instead of the largest evaluated window at *w*_*max*_ = 10, 000. This is likely explained by the observation that, beyond a certain window size, the more likely it is that the same pair of strobes is found and linked. We also looked at how the runtime scaled with window size. Figure 5 (right panel) shows the median runtime from 10 runs on the *E. coli* genome of 5.5 million nucleotides. We observe that our BST implementation greatly outperforms ℓ_*CC*_.

**Fig. 5:**
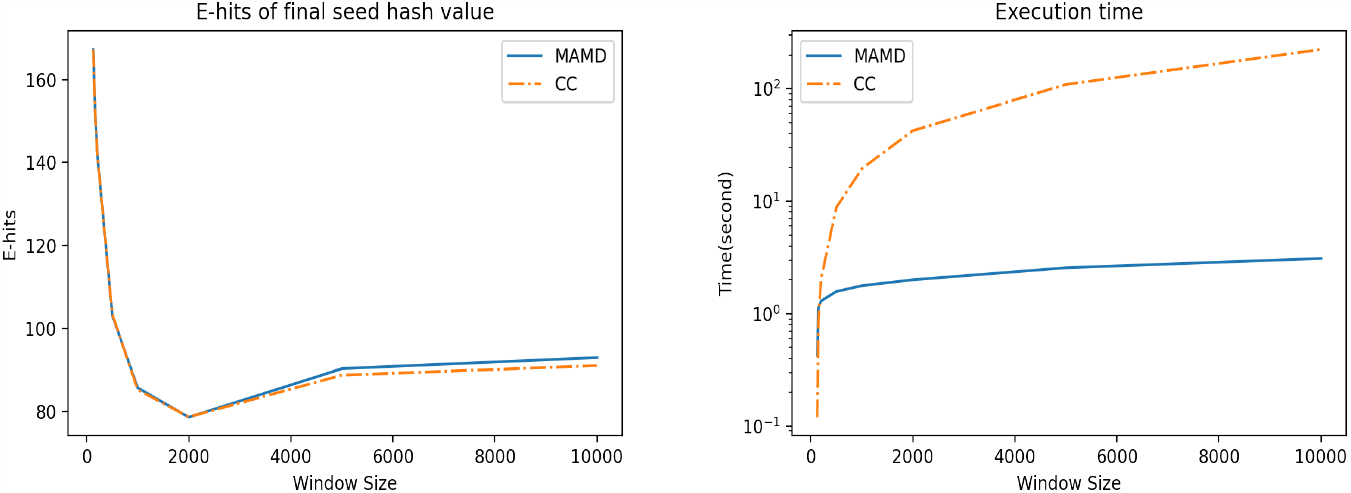
A comparison between ℓ_*MAMD*_ and ℓ_*CC*_ with parameters (*n* = 2, *l* = 128, *w*_*min*_ = 129, *w*_*max*_ = *x*), where *x* is plotted on the x-axis. Left panel shows E-hits on Chromosome Y from the CHM13 human assembly [16]. The right panel shows median runtime out of 10 runs on an *E. coli* genome of 5.5 million nt.

**Fig. 6:**
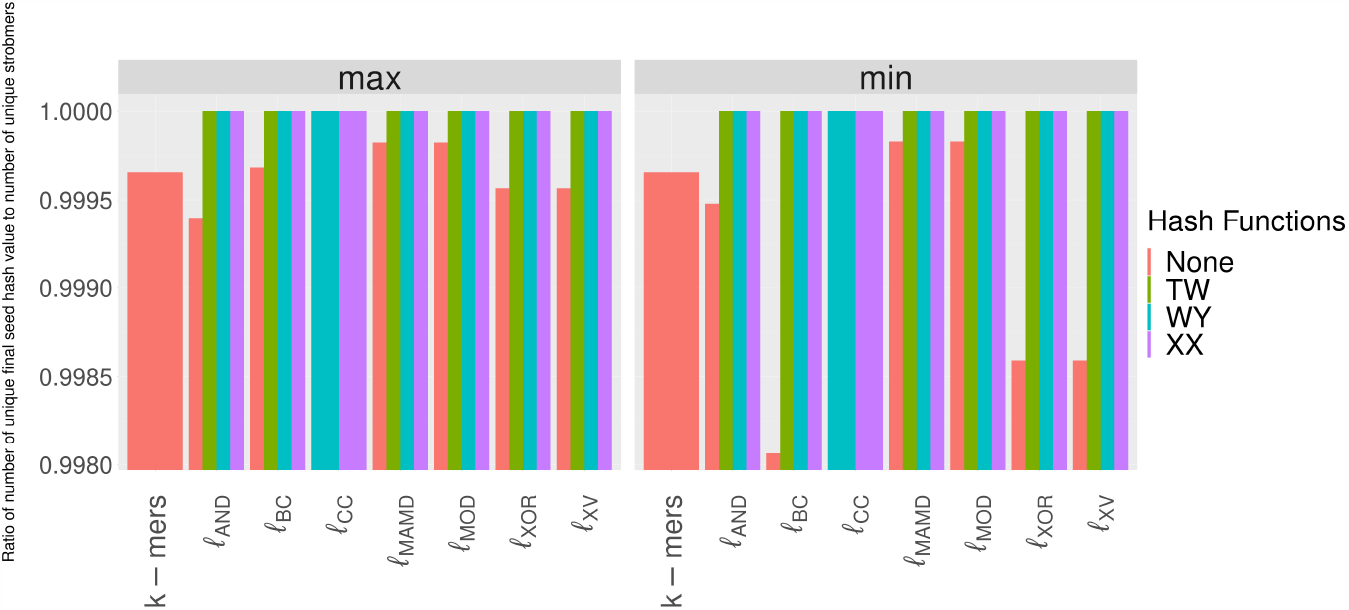
Ratio of number of unique final seed hash value to number of unique strobemers for (*n* = 2, *l* = 20, *w*_*min*_ = 21, *w*_*max*_ = 100).

### 3.5 Implementing *c*_*max*_ in strobealign

We observed in our benchmark (Fig. 2) that *c*_*min*_ together with ℓ_*BC*_ were particularly bad in terms of seed uniqueness and randomness (Fig. 2 and 3). Strobealign [20] is a short-read mapper that uses ℓ_*BC*_ together with the *c*_*min*_. Guided by our benchmark, we wanted to investigate whether *c*_*max*_ would result in better mapping results. First, strobealign adds other modifications to the strobemer constructions, such as thinning out the *k*-mers by using syncmers [6], masking the majority of bits before applying ℓ_*BC*_, and applying customized window sizes (*w*_*min*_ and *w*_*max*_) based on thinning rate and read length. Such modifications may make the observations from our analysis less effective or even inapplicable to the seeding within strobealign.

Nevertheless, we evaluated the accuracy of strobealign (v0.11.0) when mapping reads to the drosophila, CHM13, maize, and rye genomes used in [20] for read lengths 50, 75, 100, 150, 200, 250, 300, 500 by simulating one million read pairs (reads if single-end experiment) for each instance. While we did not observe a direct improvement in accuracy only when comparing the accuracy results between the two versions for neither paired-end (Table 1) nor single-end reads Table S1, we observed a large improvement in accuracy when combining the results of the two runs of strobealign. In the case of the paired-end reads (Table 1), the combined results were obtained as follows. We pick the alignment from *c*_*max*_ if the read pair was properly paired with *c*_*max*_ but not with *c*_*min*_ or the sum of alignment scores for *c*_*max*_ was higher than for *c*_*min*_ where unmapped reads count as having a score of 0. Otherwise, we picked the result from *c*_*min*_. For the single-end reads, the combined results were obtained by selecting the best alignment per read (decided by alignment score) when comparing the SAM files, where unmapped reads count as having a score of 0.

**Table 1:** Strobealign accuracy results (% of total read pairs) when mapping paired-end reads using the min comparator, max comparator, and when selecting the best read alignment between the two versions (combined). The diff column shows the percent point difference in accuracy between combined and min-comparator only, and the rightmost column indicate the difference of the combined results to that of BWA, the overall most accurate aligner in the benchmark in [20]. Negative values indicate that strobealign has higher accuracy.

| dataset (genome-read length) | min comparator | max comparator | combined | diff | combined diff to BWA |
| --- | --- | --- | --- | --- | --- |
| drosophila-50 | 90.1890 | 90.1542 | 90.4216 | +0.2326 | +0.0901 |
| drosophila-75 | 91.6441 | 91.6422 | 91.6818 | +0.0377 | -0.0233 |
| drosophila-100 | 92.3879 | 92.3948 | 92.4168 | +0.0289 | -0.0207 |
| drosophila-150 | 93.2112 | 93.2220 | 93.2302 | +0.0190 | -0.0350 |
| drosophila-200 | 93.5209 | 93.5309 | 93.5347 | +0.0138 | -0.0617 |
| drosophila-300 | 95.3608 | 95.3681 | 95.3734 | +0.0126 | -0.0425 |
| drosophila-500 | 95.6936 | 95.7124 | 95.7132 | +0.0196 | -0.0774 |
| CHM13-50 | 90.6350 | 90.5854 | 91.0971 | +0.4621 | +0.5227 |
| CHM13-75 | 92.5158 | 92.5197 | 92.6462 | +0.1304 | +0.2038 |
| CHM13-100 | 93.2198 | 93.2153 | 93.3182 | +0.0983 | +0.1367 |
| CHM13-150 | 94.1404 | 94.1486 | 94.2101 | +0.0698 | +0.0543 |
| CHM13-200 | 94.4340 | 94.4397 | 94.4870 | +0.0530 | +0.0241 |
| CHM13-300 | 95.6266 | 95.6271 | 95.6779 | +0.0512 | +0.0796 |
| CHM13-500 | 95.9505 | 95.9555 | 96.0153 | +0.0648 | +0.0523 |
| rye-50 | 69.1402 | 68.9105 | 71.1016 | +1.9613 | +2.3892 |
| rye-75 | 80.5345 | 80.4464 | 81.5855 | +1.0511 | +1.4724 |
| rye-100 | 85.6483 | 85.6312 | 86.4098 | +0.7615 | +0.9966 |
| rye-150 | 90.2038 | 90.2065 | 90.6332 | +0.4295 | +0.4415 |
| rye-200 | 91.4773 | 91.4661 | 91.7506 | +0.2733 | +0.1812 |
| rye-300 | 94.5574 | 94.5816 | 94.6644 | +0.1070 | +0.1012 |
| rye-500 | 95.1326 | 95.1618 | 95.2114 | +0.0787 | +0.0374 |
| maize-50 | 71.4703 | 71.3223 | 73.0630 | +1.5927 | +1.6149 |
| maize-75 | 82.1255 | 82.0405 | 82.9049 | +0.7794 | +0.7763 |
| maize-100 | 87.1317 | 87.1404 | 87.7111 | +0.5793 | +0.5152 |
| maize-150 | 91.6731 | 91.6841 | 91.9923 | +0.3191 | +0.1784 |
| maize-200 | 92.9204 | 92.9328 | 93.1210 | +0.2005 | +0.0883 |
| maize-300 | 96.7084 | 96.7183 | 96.8246 | +0.1163 | +0.0332 |
| maize-500 | 97.2899 | 97.2962 | 97.4021 | +0.1122 | -0.0025 |

We observed that both in the paired-end and single-end experiments, the shorter read lengths benefited the most from combining the results (seen from the percentage point difference in Table 1 and S1. The generally most accurate aligner in the benchmark of strobealign [20] was BWA-MEM [9]. We included the percentage point difference to BWA-MEM in Table 1 and S1, where a negative result indicates that the accuracy from the combined experiment was more accurate than BWA-MEM. While strobealign is more accurate when aligning reads to drosophila for most read lengths, BWA-MEM is more accurate on the larger genomes CHM13, rye, and maize. However, by comparing the diff column to the combined diff to BWA column, the percentage point difference to BWA-MEM is decreased by half or more for many of the datasets, when combined results are used.

For the single-end experiment, we also generated a combined version where we only used the *c*_*max*_ result if a read was unmapped with the default strobealign (i.e., using *c*_*min*_). This version also generated a relatively large increase in accuracy, particularly for the shorter reads (Table S2). This suggests that expensive alignment rescue steps (both in the seeding step and in the alignment step) may be avoided for the shorter reads by having more matching seeds.

In addition, we observed no apparent difference in the number of mapped reads, memory usage, and runtime between the two versions of strobealign. Our results suggest a mapping strategy where the *c*_*min*_ and *c*_*max*_ comparators could be combined to allow for more accurate read alignment with strobealign for the shortest read lengths. While the combined results were obtained as a proof-of-concept by running strobealign twice, more efficient solutions could be implemented, as discussed in future work.

## 4 Discussion and conclusions

Constructing randstrobes can be split into four modular operations: computing a hash value of the individual strobes (hash function), computing a hash value of two linked strobes (link function), selecting the final randstrobe out of several candidates through a comparator function, and computing the final randstrobe hash value of the selected randstrobe. The three first operations (hash, link, and comparator functions) produce different results depending on their implementation. We proposed and evaluated new hash link and comparator methods to construct randstrobes. We also designed metrics for evaluating the bias of different methods. Our evaluation metrics and benchmark across several different combinations of operations to produce randstrobes uncovered biases and limitations in previously proposed techniques. Our evaluation provides general guidelines for which method to use in the three steps when considering using randstrobes as seeds for sequence comparison applications. From our evaluation we conclude the following.

- **Hashing:** Always use a hash function to hash the strobes before linking. It does not result in a large overhead in construction time while being beneficial for pseudo-randomness for most link functions. The hash functions has roughly the same pseudo-randomness performance, but *h*_*WY*_ function had the best runtime performance for the expensive-methods class in our evaluation.
- **Comparator:** For repetitive sequences with occasional variations, such as in the dataset we benchmarked on, a max comparator will be inclined to select strobes that are different. In contrast, if present, the min comparator will select identical strobes for some of the link functions, resulting in lower uniqueness if several repeated copies are used. Since we did not observe any notable difference in computation time between the min and max comparators, we suggest always using the max comparator when implementing randstrobes.
- **Link function:** There is a trade-off between execution time and pseudo-randomness performance. The slower ℓ_*CC*_ and ℓ_*XV*_ has the highest pseudo-randomness (Fig. 2), but are more expensive to compute (Fig. 4). The recommended method would depend on the needs of the application and the window size.
- **Pitfalls:** First, the ℓ_*MOD*_ function should be used together with *c*_*min*_ due to its selection of identical strobes in repetitive regions which causes excessive repetitiveness of seeds. Second, the ℓ_*MOD*_ does not offer beneficial pseudo-randomness and is computationally more expensive compared to other methods in the same class. Therefore, we do not recommend its use in any scenario.
- **Large windows:** The construction time of randstrobes depends on the window size, and they can become computationally expensive to compute for large windows. The ℓ_*MAMD*_ link function can over-come the high construction time and scales well for very large windows. However, it has slightly higher repetitiveness compared to the expensive class methods (Fig. 5).

We also observed how results in the bioinformatic tool strobealign, using strobemer-based seeding, produce different results under different implementations. Suggesting putting further thought into the underlying method for constructing randstrobes.

### 4.1 Future work

#### Using a rolling hash function

The cheap computation methods separate the steps of applying a hash function to the strobes from applying the link function. This class of methods may benefit from using a rolling hash function, such as the one proposed in [14], that can be applied to the hash computation. Such optimization is beneficial primarily if the hashing is expensive relative to the linking, which is not the case for lager window sizes. However, by arithmetic reasoning, we designed a link function ℓ_MAMD_ that reduced the time complexity of the construction. It remains to be investigated whether the rolling hash approach allows for arithmetic operations that could reduce the computations in the linking step.

#### Combining min and max comparators

We found improved accuracy when combining results from the min and max comparators in strobealign. Our proof-of-concept approach involved running strobealign twice and post-processing the alignments, resulting in slightly more than twice the runtime compared to a single run. To mitigate the runtime and memory doubling, integrating seeds from both comparators into strobealign is a solution. This would only double the vector containing the seeds in memory; for instance, it would increase from 9.6GB to 19.2GB for CHM13 data. This memory increase leads to a peak memory in strobealign rising from 14.7GB to 24.3GB. It is not obvious that the runtime would increase over current strobealign. Most datasets have uniquely mapped reads sharing the same candidate locations from both comparators. Also, extension alignment, a bottleneck in strobealign, can be done once for most reads, and costly alignment rescue steps, performed especially for shorter reads, would be reduced in a combined minmax version. One could consider implementing such a high sensitivity setting within strobealign for the shortest read lengths, compromising memory usage.

## 5 Data availability

Our scripts to generate data, construct randstrobes, and perform the benchmarks of the methods are found at https://github.com/Moein-Karami/RandStrobes. The scripts for running the strobealign analysis are found at https://github.com/marcelm/K_Sahlin_2201/tree/main.

## 7 Acknowledgements

We thank Daniel Liu for suggesting the ℓ_*CC*_ link function and Heng Li for useful feedback on the linking methods.

## A Appendix

### A.1 ℓ_MAMD_ implementation

Link functions ℓ_MOD_ and ℓ_MAMD_ use the same modulo operation. However, ℓ_MAMD_ reduces the time complexity of the construction through the use of a Binary Search Tree (BST). The ℓ_MAMD_ method is implemented as follows. Consider the min comparator and a BST *B* containing all the possible candidates 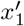 for the second strobe *x*_1_ by storing the hash of each strobe modulo *p*. We want to choose 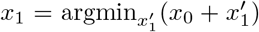 mod *p*. First note that we have 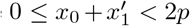 for all possible strobes 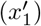 in the window, because we store all the hash values modulo *p*. There are two possibilities for the best value 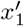 that we discuss separately.

- If 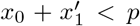, the best possible candidate is the smallest value in the window. This is because for any other value 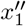 that is greater than 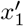 and satisfies 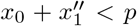, we have 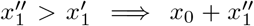 mod 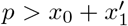 mod *p*.
- If 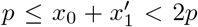, the best possible candidate is the smallest value that is equal to or greater than *p* − *x*_0_ in the window. This is because for any other value 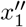 that is greater than 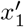, we have 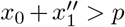 and 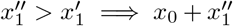 mod 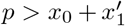 mod *p*. For any other value 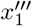 that is smaller than 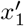, we have 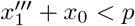, which falls under the previous situation.

Therefore, the only two candidates for 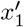 are the minimum value in *B* and the smallest value that is equal to or greater than *p* − *x*_0_. We can compare the two values and select the best one. These values can both be found in *O*(log(*W*_max_ − *W*_min_)) time, where *W*_max_ and *W*_min_ are the boundaries of the window. Finding the minimum element in a BST is a standard operation. For the second case, in the BST implementation we use (std::set in C++), we can find the greatest value *y ≤ p* − *x*_0_. We can then find the next element in *O*(1) which implies the smallest value that is greater than *p* − *x*_0_.

To create the next randstrobe, the window swaps the value corresponding to the leftmost value in the window with an incoming value (rightmost value in the new window). Removing and adding values are also *O*(log(*W*_max_ − *W*_min_)) operations in a BST. If a max comparator is used, we have an analogous case.

### A.2 Results for randstrobes with three strobes

We also investigated constructing randstrobes with (*n* = 3, *l* = 20, *w*_*min*_ = 21, *w*_*max*_ = 100). In general, we observed similar behavior in terms of pseudo-randomness as for the construction of randstrobes with two strobes (Fig. 7). This is expected due to the recursive nature of the construction. That is, when selecting the *m*-th strobe, we perform the selection based on a base value constructed from previous strobes (described in section 2.3). This process repeats recursively. When *m* = 2, the base value is simply the hash value of the first strobe.

**Fig. 7:**
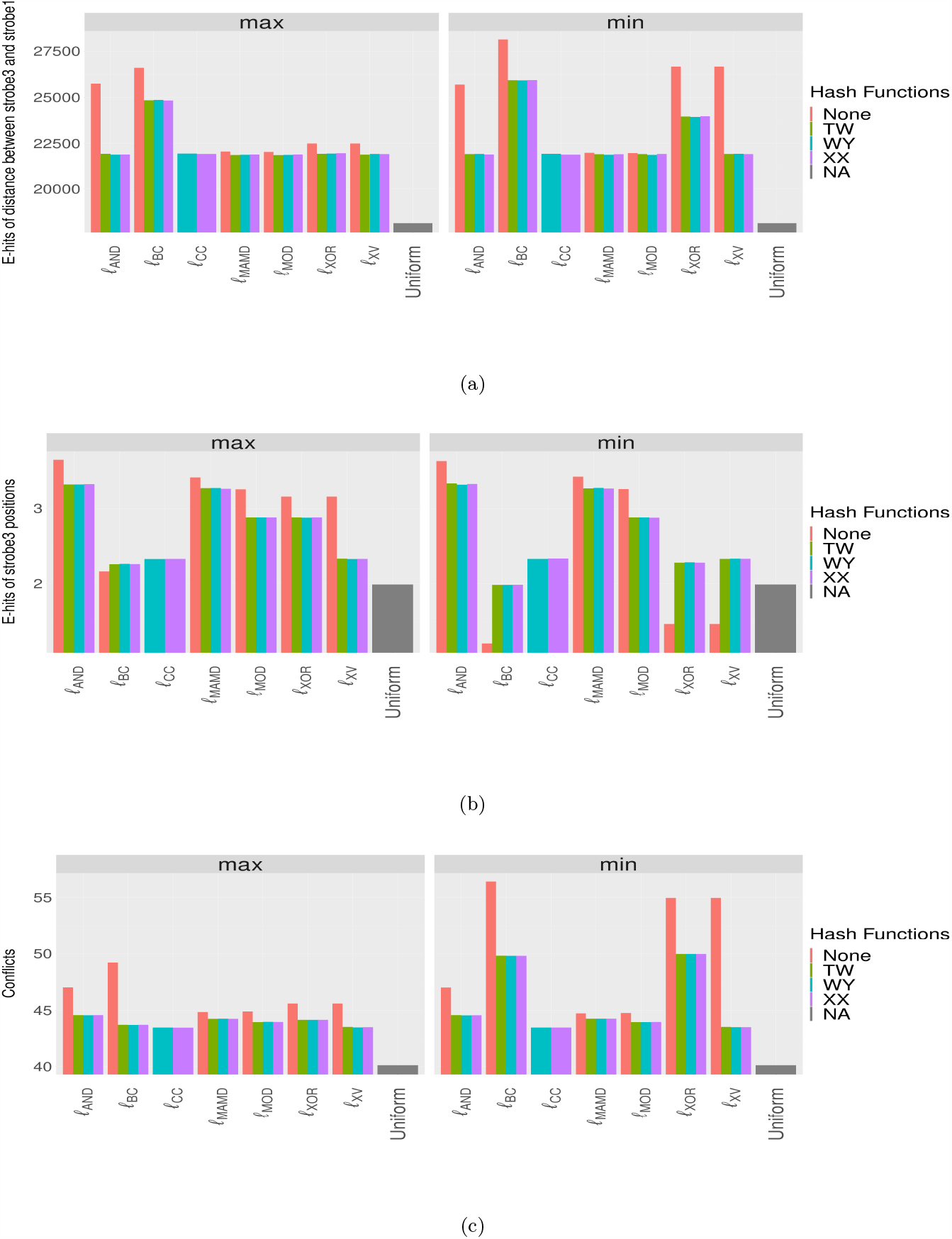
Results for metrics *E*_*d*_, *E*_*p*3_, and *C* for randstrobes with parameter settings (*n* = 3, *l* = 20, *w*_*min*_ = 21, *w*_*max*_ = 100) for the repetitive sequence dataset. The x-axis shows the different linking methods and the reference value denoted uniform, indicating near perfect randomness. For metrics *E*_*d*_ and *C*, a low value is desirable. For *E*_*p*3_, a value close to uniform is desirable, and is not necessarily as low as possible, due to bias B in Fig. 1. The results for the max and min comparator are shown in left and right panels in each subfigure, respectively.

As for the uniqueness of seeds, as demonstrated in [18], randstrobes with three strobes are relatively more unique than *k*-mers with the same number of sampled bases (Fig. 8, *k* = 60). This is due to the increase in range of the seed, thus, the increased number of options available for selecting strobes. But some combinations (e.g., no hashing or based on the XOR operator; Fig. 8) can reduce the uniqueness.

**Fig. 8:**
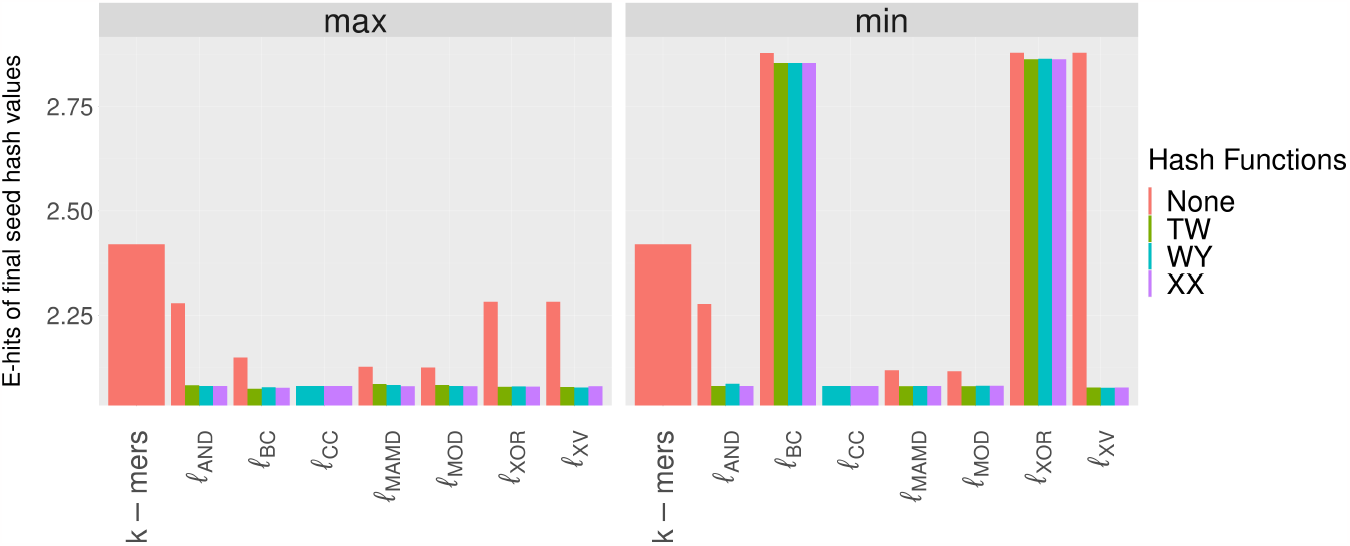
E-hits of final seed hash values for randstrobes with three strobes with parameter settings (*n* = 3, *l* = 20, *w*_*min*_ = 21, *w*_*max*_ = 100) compared to *k*-mers with *k* = 60

### A.3 Figures

### A.4 Tables

**Table S1:** Strobealign accuracy results (% of total reads) when mapping single-end reads using the min comparator, max comparator, and when selecting the best read alignment between the two versions (combined). The diff column shows the difference between combined and min comparator only, and the rightmost column indicate the difference of the combined results to that of BWA, the overall most accurate aligner in the benchmark in [CIT strobealign].

| dataset (genome-read length) | min comparator | max comparator | combined | diff | combined diff to BWA |
| --- | --- | --- | --- | --- | --- |
| drosophila-50 | 82.5118 | 82.3825 | 84.2688 | +1.7570 | +2.5606 |
| drosophila-75 | 87.9944 | 87.9556 | 88.2902 | +0.2958 | +0.2191 |
| drosophila-100 | 89.2585 | 89.2853 | 89.4715 | +0.2130 | +0.1209 |
| drosophila-150 | 90.9438 | 90.9455 | 91.0235 | +0.0797 | +0.0184 |
| drosophila-200 | 91.9501 | 91.9644 | 91.9927 | +0.0426 | +0.0286 |
| drosophila-300 | 93.2169 | 93.2451 | 93.2396 | +0.0227 | +0.0067 |
| drosophila-500 | 94.5768 | 94.5895 | 94.6109 | +0.0341 | +0.0162 |
| CHM13-50 | 81.6910 | 81.5544 | 83.6503 | +1.9593 | +2.9080 |
| CHM13-75 | 88.9220 | 88.9050 | 89.4662 | +0.5442 | +0.6695 |
| CHM13-100 | 90.6392 | 90.6112 | 91.0038 | +0.3646 | +0.3714 |
| CHM13-150 | 92.3990 | 92.3885 | 92.5668 | +0.1678 | +0.1467 |
| CHM13-200 | 93.2258 | 93.2383 | 93.3355 | +0.1097 | +0.0965 |
| CHM13-300 | 94.1503 | 94.1331 | 94.2176 | +0.0673 | +0.1184 |
| CHM13-500 | 95.0598 | 95.0737 | 95.1460 | +0.0862 | +0.1218 |
| rye-50 | 44.6869 | 44.4947 | 46.1874 | +1.5005 | +2.1252 |
| rye-75 | 60.2210 | 60.0863 | 61.3118 | +1.0908 | +1.5198 |
| rye-100 | 69.3358 | 69.3736 | 70.4965 | +1.1607 | +1.5197 |
| rye-150 | 80.4388 | 80.3968 | 81.3871 | +0.9483 | +0.8698 |
| rye-200 | 85.9181 | 85.8857 | 86.6284 | +0.7103 | +0.6255 |
| rye-300 | 90.5230 | 90.5150 | 90.9841 | +0.4611 | +0.5704 |
| rye-500 | 93.5070 | 93.5344 | 93.7914 | +0.2844 | +0.3686 |
| maize-50 | 47.4174 | 47.2713 | 48.8452 | +1.4278 | +2.0588 |
| maize-75 | 61.9111 | 61.8251 | 62.8084 | +0.8973 | +1.1361 |
| maize-100 | 70.5000 | 70.4634 | 71.4263 | +0.9263 | +1.0928 |
| maize-150 | 81.1708 | 81.1807 | 81.9200 | +0.7492 | +0.5582 |
| maize-200 | 86.7010 | 86.7061 | 87.2567 | +0.5557 | +0.3617 |
| maize-300 | 91.8729 | 91.8704 | 92.2597 | +0.3868 | +0.3108 |
| maize-500 | 95.4025 | 95.4229 | 95.7271 | +0.3246 | +0.2333 |

**Table S2:** Strobealign accuracy results (% of total reads) when mapping single-end reads using the min comparator, max comparator, and when adding the reads that were mapped only with the max comparator to the mapping results with the min comparator (combined). The diff column shows the difference between combined and min comparator only.

| dataset (genome-read length) | min comparator | max comparator | combined | diff |
| --- | --- | --- | --- | --- |
| drosophila-50 | 82.5118 | 82.3825 | 84.1942 | +1.6824 |
| drosophila-75 | 87.9944 | 87.9556 | 88.2319 | +0.2375 |
| drosophila-100 | 89.2585 | 89.2853 | 89.4176 | +0.1591 |
| drosophila-150 | 90.9438 | 90.9455 | 90.9823 | +0.0385 |
| drosophila-200 | 91.9501 | 91.9644 | 91.9590 | +0.0089 |
| drosophila-300 | 93.2169 | 93.2451 | 93.2170 | +0.0001 |
| drosophila-500 | 94.5768 | 94.5895 | 94.5769 | +0.0001 |
| CHM13-50 | 81.6910 | 81.5544 | 83.1962 | +1.5052 |
| CHM13-75 | 88.9220 | 88.9050 | 89.1354 | +0.2134 |
| CHM13-100 | 90.6392 | 90.6112 | 90.7793 | +0.1401 |
| CHM13-150 | 92.3990 | 92.3885 | 92.4377 | +0.0387 |
| CHM13-200 | 93.2258 | 93.2383 | 93.2328 | +0.0070 |
| CHM13-300 | 94.1503 | 94.1331 | 94.1504 | +0.0001 |
| CHM13-500 | 95.0598 | 95.0737 | 95.0598 | +0.0000 |
| rye-50 | 44.6869 | 44.4947 | 45.2202 | +0.5333 |
| rye-75 | 60.2210 | 60.0863 | 60.2857 | +0.0647 |
| rye-100 | 69.3358 | 69.3736 | 69.3846 | +0.0488 |
| rye-150 | 80.4388 | 80.3968 | 80.4509 | +0.0121 |
| rye-200 | 85.9181 | 85.8857 | 85.9201 | +0.0020 |
| rye-300 | 90.5230 | 90.5150 | 90.5230 | +0.0000 |
| rye-500 | 93.5070 | 93.5344 | 93.5070 | +0.0000 |
| maize-50 | 47.4174 | 47.2713 | 48.0640 | +0.6466 |
| maize-75 | 61.9111 | 61.8251 | 61.9915 | +0.0804 |
| maize-100 | 70.5000 | 70.4634 | 70.5609 | +0.0609 |
| maize-150 | 81.1708 | 81.1807 | 81.1898 | +0.0190 |
| maize-200 | 86.7010 | 86.7061 | 86.7042 | +0.0032 |
| maize-300 | 91.8729 | 91.8704 | 91.8729 | +0.0000 |
| maize-500 | 95.4025 | 95.4229 | 95.4025 | +0.0000 |

